# RACIPE: A computational tool for Modeling Gene Regulatory Circuits using Randomization

**DOI:** 10.1101/210419

**Authors:** Bin Huang, Dongya Jia, Jingchen Feng, Herbert Levine, José N. Onuchic, Mingyang Lu

## Abstract

**Motivation:** One of the major challenges in traditional mathematical modeling of gene regulatory circuits is the insufficient knowledge of kinetic parameters. These parameters are often inferred from existing experimental data and/or educated guesses, which can be time-consuming and error-prone, especially for large networks.

**Results:** We present a computational tool based on our newly developed method named <u>*ra*</u>ndom <u>*ci*</u>rcuit <u>*pe*</u>rturbation (RACIPE), to explore the robust dynamical features of gene regulatory circuits without the requirement of detailed kinetic parameters. RACIPE generates an ensemble of circuit models with distinct random parameters and uniquely identifies robust dynamical properties by statistical analysis. Here, we discuss software implementation and illustrate the usage of RACIPE on coupled toggle-switch circuits and a published circuit of B-lymphopoiesis. We expect RACIPE to contribute to a more comprehensive and unbiased understanding of gene regulatory mechanisms.

**Availability:** RACIPE is a free open source software distributed under (Apache 2.0) license and can be downloaded from GitHub (https://github.com/simonhb1990/RACIPE-1.0).

## 1 Introduction

Biological processes are orchestrated by complex gene regulatory networks (GRNs). To understand the operating principles of GRNs, mathematical modeling approaches (Smolen *et al.*, 2000; Novère, 2015) have been widely used in various contexts, such as regulation of cell cycle (Tyson, 1991), stem cell development (Huang *et al.*, 2007), circadian rhythm (Smolen *et al.*, 2001), developmental pattern formation (Reeves *et al.*, 2006) and cell phenotypic switches in cancer (Ao *et al.*, 2008; Lu *et al.*, 2013; Zhang *et al.*, 2014; Li and Wang, 2015; Yu *et al.*, 2017). To model the dynamics of GRNs, different computational algorithms have been developed (Dehmer *et al.*, 2011), such as ordinary differential equations (ODEs)-based models (Strogatz, 2007), Boolean network models (Li *et al.*, 2004; Steinway *et al.*, 2014), Bayesian network models (Zhang *et al.*, 2013), agent-based models (Robertson-Tessi *et al.*, 2015), and reaction-diffusion models (Roberts *et al.*, 2013). The ODEs-based models consider more regulatory details compared to Boolean or Bayesian network models and less computationally intensive than agent-based model and reaction-diffusion models, thus being a very attractive approach to simulate the operation of GRNs.

It is believed that there is a core gene regulatory circuit underlying a GRN which functions as a decision-making module for one specific biological process (Milo *et al.*, 2002; Hartwell *et al.*, 1999). Identification of such core gene circuits can largely reduce the complexity of network modeling. Notably, the core gene regulatory circuit doesn’t function isolatedly. Instead, its operation is usually regulated by other genes and signaling pathways (“peripheral factors”) that interact with the core circuit. Although the ODE-based and other modeling approach have been successfully applied to analyze the dynamics of the core gene circuits in certain scenarios, these approaches typically suffer from two issues. First, it is very difficult for traditional modeling approach to consider the effects of these “peripheral” factors due to their inherent complexity. Second, the modeling approaches are usually limited by insufficient knowledge of the kinetic parameters for many of the biological processes. In this case, the value of most parameters have to be inferred either by educated guess or fitting to the experimental results, which can be time-consuming and error-prone especially for large gene networks.

To deal with these issues, we previously established a new computational method, named <u>*ra*</u>ndom <u>*ci*</u>rcuit <u>*pe*</u>rturbation (RACIPE), to study the robust dynamical features of gene regulatory circuits without the requirement of detailed kinetic parameters (Huang *et al.*, 2017). RACIPE takes the topology of the core regulatory circuit as the *only* input and unbiasedly generates an ensemble of mathematical models, each of which is characterized by a unique set of kinetic parameters. For each mathematical model, it contains a set of chemical rate equations, which are subjected to non-linear dynamics analysis. From the ensemble of models, we can analyze the robust dynamical properties of the core circuit by statistical analysis. In RACIPE, the effects of the “peripheral factors” are modeling as random perturbations to the kinetic parameters. Unlike the traditional ODEs-based modeling, RACIPE uses a self-consistent scheme to randomize all kinetic parameters for each mathematical model instead of relying on a particular set of parameters. Unlike other methods using randomization (Meir *et al.*, 2002; Feng *et al.*, 2004; Gutenkunst *et al.*, 2007; Llamosi *et al.*, 2016), RACIPE adopts a more carefully designed sampling strategy to randomize parameters across a wide range while satisfying the half-function rule, where each regulatory link has about 50% chance to be activated in the ensemble of RACIPE models. RACIPE-generated gene expression data and corresponding parameters can be analyzed by statistical learning methods, such as hierarchical clustering analysis (HCA) and principle component analysis (PCA) which provides a holistic view of the dynamical behaviors of the gene circuits. Notably, RACIPE integrates statistical learning methods with parameter perturbations, which makes it distinct from the traditional parameter sensitivity analysis (Meir *et al.*, 2002; Feng *et al.*, 2004), parameter space estimation (Leon *et al.*, 2016) and other randomization strategies (Gutenkunst *et al.*, 2007; Llamosi *et al.*, 2016). In addition, our previous work shows that robust gene expression patterns are conserved against large parameter perturbations due to the restraints from the circuit topology. Thus we can interrogate the dynamical property of a gene circuit by randomization.

Without the need to know detailed kinetic parameters, RACIPE can 1) identify conserved dynamical features of a relatively large gene regulatory circuits across an ensemble of mathematical models; and 2) generate predictions on gain-of-function and loss-of-function mutations of each gene/regulatory link; and 3) discover novel strategies to perturb particular cell phenotypes. The application of RACIPE to a proposed core 22-gene regulatory circuit governing epithelial-to-mesenchymal transition (EMT) showed that RACIPE captures experimentally observed stable cell phenotypes, and predict the efficiency of various biomarkers in distinguishing different EMT phenotypes (Huang *et al.*, 2017).

Here, we report a computational tool that we developed based on the random circuit perturbation method. In the following, we first discuss the implementation of RACIPE, including how the tool processes the input topology file of a gene network, estimates the range of parameters for randomization and solves stable steady states, *etc*. By applying RACIPE on a coupled toggle-switch circuit, we evaluate the computational cost of using RACIPE, detail the procedure on how to choose an appropriate number of RACIPE models and number of initial conditions for each RACIPE model to get converged simulation results for a gene circuit, and further illustrate how to do perturbation analysis using RACIPE. Lastly, we apply RACIPE on a published gene circuit governing B-lymphopoiesis (Salerno *et al.*, 2015) and show that RACIPE can capture multiple gene expression states during B cell development and the fold-change in expression of several key regulators between stages (van Zelm *et al.*, 2005). In summary, we expect RACIPE to be a valuable tool to decipher the robust dynamical features of gene circuits in many applications.

## 2 Methods

### 2.1 Overview of Random Circuit Perturbation

Random Circuit Perturbation (RACIPE) method is developed to identify the robust dynamical features of a biological gene circuit without the need of detailed circuit parameters (Huang *et al.*, 2017). The method is composed of two parts (Fig. 1a) – a procedure to generate and simulate an ensemble of models and statistical analysis across all generated models to identify robust dynamical features of the circuit.

**Fig. 1.**
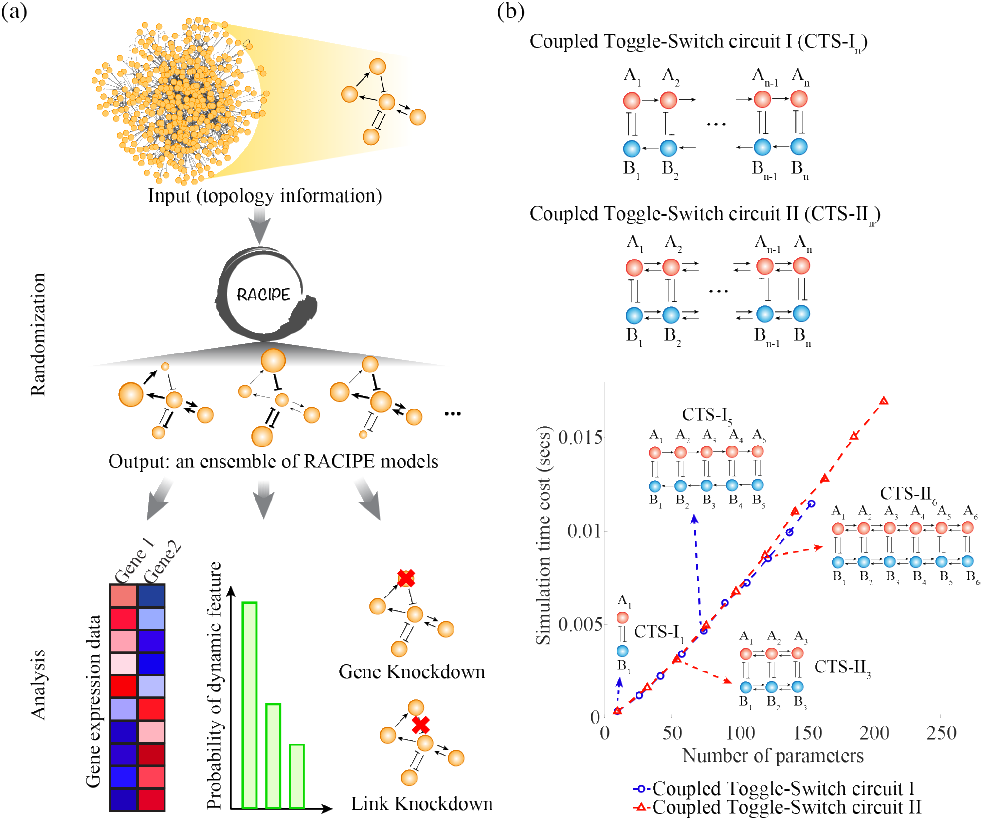
The method of random circuit perturbation (a) Workflow of RACIPE. It first generates an ensemble of models by randomizing the kinetic parameters of the chemical rate equations of the core circuit, and then analyzes the ensemble of models by statistical analysis. (b) RACIPE is tested on two types of coupled toggle-switch (CTS) circuits (diagram illustrated in the top panel). The arrows represent transcriptional activation; the bar-headed arrows represent transcriptional inhibition. For both cases, the average time cost to simulate a RACIPE model (y-axis) is linearly proportional to the number of model parameters (x-axis).

### 2.2 Software Implementation

We report a tool based on the RACIPE method specifically for multi-stable gene regulatory circuits. With the input of the topology of a gene circuit, the tool automatically builds the mathematical model of the circuit, randomizes the model parameters, and solves the model to output the solutions of the stable steady states. These results can be used to uncover the robust features of the circuit, such as the stable steady-state gene expressions. The main steps of the tool are elaborated below.

#### 2.2.1 Input data

The main input of RACIPE is the topology of a gene circuit, i.e. the gene names and the regulatory links connecting them. The current version can be applied to gene regulatory circuits with only transcription factors. We will expand its capacity to other regulation types in the future. In the input topology file (e.g., “circuit.topo”), each line specifies a regulatory link, which contains the name of source gene, the name of target gene, and the type of interactions (activation or inhibition). The list of gene nodes is not required, as it is automatically generated in RACIPE. Table 1 shows an example of the input topology file for a toggle-switch circuit, which has two mutually inhibiting genes A and B.

**Table.1.** Format of the input topology file (“circuit.topo”)

| Source | Target | Type<sup>#</sup> |
| --- | --- | --- |
| A | B | 2 |
| B | A | 2 |

<sup>#</sup> “1” stands for activation, and “2” stands for inhibition.

#### 2.2.2 Process Circuit Topology Information

Based on the input circuit topology, RACIPE automatically builds mathematical models of ordinary differential equations (ODEs). For instance, the temporal dynamics of a toggle switch circuit can be modeled by the following ODEs: 
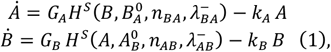
where *A* and *B* are the expression levels of genes A and B, respectively. *G*_*A*_ and *G*_*B*_ are the maximum production rates (the production rate for the gene with all activators, but not any inhibitor, binding to the promoter). *k*_*A*_ and *k*_*B*_ are the innate degradation rates of A and B, respectively. The effects of the inhibitory regulation of gene A by B is formulated as a non-linear shifted Hill function (Lu *et al,* 2013) 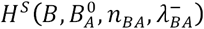 defined as 
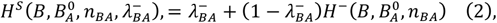
where 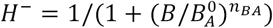 is the inhibitory Hill function, 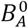 is the threshold level, *n*_*BA*_ is the Hill coefficient and 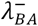 is the maximum fold change of the B level caused by the inhibitor A 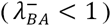. The inhibition of gene A by gene B can be modeled in a similar way.

When multiple regulators target a gene, the functional form of the rate equations depends on the nature of the multivalent regulation. Currently, we adopt a common scheme where we assume that these regulatory interactions are independent. Thus, the overall production rate is written as the product of the innate production rate of the target gene and the shifted Hill functions for all the regulatory links. We will consider other cases, such as competitive regulation, in a later version.

#### 2.2.3 Estimate the Ranges of Parameters for Randomization

Next, RACIPE estimates, for each parameter, the range of values for randomization. Most of the parameter ranges, such as the ones for production and degradation rates, are preset (see SI 1.1), while the ranges for the threshold values from the shift Hill functions are estimated numerically to satisfy the “half-functional” rule. The “half-functional” rule ensures that each link in the circuit has roughly 50% chance to be functional across all the models (Huang *et al.,* 2017). All the parameter ranges are generated and stored in a parameter file (“circuit.prs”).

#### 2.2.4 Solve and Identify the Stable Steady States

To generate a model, RACIPE randomizes each parameter independently within the pre-calculated range. For each model with a particular set of parameters, RACIPE numerically simulates the dynamics of the model (see SI 1.2). To identify all possible stable steady states of each model, we repeat the simulations for multiple times with different initial conditions, randomly chosen from a log-uniform distribution ranging from the minimum possible level to the maximum possible level. To rapidly obtain the stable steady states, we simulate the dynamics by the Euler method with a large time step. From the steady state solutions of all the realizations, we identify distinct stable states, defined as those whose Euclidean distances of the levels among them are all larger than a small threshold (see SI 1.3). The above procedure is repeated for all the models. Together, we obtain a large set of gene expression data and model parameters for statistical analysis. In the implementation, RACIPE generates nRM number of random models, each of which is subject to simulations from nIC number of initial conditions. We will discuss how to appropriately choose nRM and nIC in the Results section.

#### 2.2.5 Output Data

Lastly, the model parameters and the steady state gene expressions of all RACIPE models are stored separately. The parameters for each RACIPE model are stored in “circuit_parameter.dat”, where each row corresponds to one RACIPE model, and each column shows the value of a parameter. The parameters follow the same order in the “circuit.prs” file. Depending on the number of stable states of a RACIPE model, its gene expressions are stored in the “circuit_solution_i.dat”, where i is the number of stable states. In the “circuit_solution_i.dat”, each row shows the gene expression vectors of all the stable steady states from a RACIPE model. These data are subject to further statistical analysis.

#### 2.2.6 Options

RACIPE allows adjusting simulation parameters by directly using them in the command line or changing them in “circuit.cfg” file (see the README file for detailed instructions). Moreover, RACIPE also has options to perform simulations of perturbations, such as gene knockout, over-expression, and removal of a regulatory link. Unlike conventional approach, RACIPE applies perturbations (see SI 1.4) to the entire ensemble of models to capture the conserved behavior of the treatment.

## 3 Results

### 3.1 Time Cost of Simulations

To evaluate the performance of RACIPE with different choices of simulation parameters, we test the tool on two types of coupled toggle-switch (CTS) circuits (Fig. 1b, see SI section 2 for mathematical models). They both contain several toggle-switch motifs, but different connecting patterns among these motifs, where the type I circuits (CTS-I) have unidirectional activations among A genes (B genes), while the type II circuit (CTS-II) have mutual activations among A genes (B genes). These circuits have been actively studied to understand the coupled cellular decision-making processes (Jolly *et al.*, 2015; Huang *et al.*, 2015). By changing the number of toggle-switch motifs, we can easily test RACIPE on circuits of different sizes. For each circuit, we generate 10,000 random models and solve steady-state expressions starting from 1000 initial conditions for each model. As shown in Fig 1b, for both types of circuits, the average simulation time to solve a RACIPE model scales linearly with the total number of parameters in the model, suggesting its potential use on large circuits. Of note, the total time to simulate all RACIPE models depends on other factors (the number of models, the number of initial conditions, etc.), which will be discussed in the next section.

### 3.2 Convergence Test

As mentioned above, there are two important simulation parameters - the number of RACIPE models (nRM) and, for each model, the number of initial conditions (nIC) that are used to find all possible stable steady states. When nRM and nIC are too small, the results from the ensemble of models may not converge and be statistically significant. However, having too large nRM and nIC sacrifices computational efficiency.

To identify an optimal choice of nRM and nIC, we test the effects of both on the convergence of the simulation results by calculating the dissimilarity of the probability distribution of the number of stable states (referred to as the “dissimilarity of states”) and the distribution of gene expressions (referred to as the “dissimilarity of expressions”) using different values of nRM and nIC. (Fig. 2 and Fig. 3). If the simulation result converges well, the dissimilarity values are expected to be small.

**Fig. 2.**
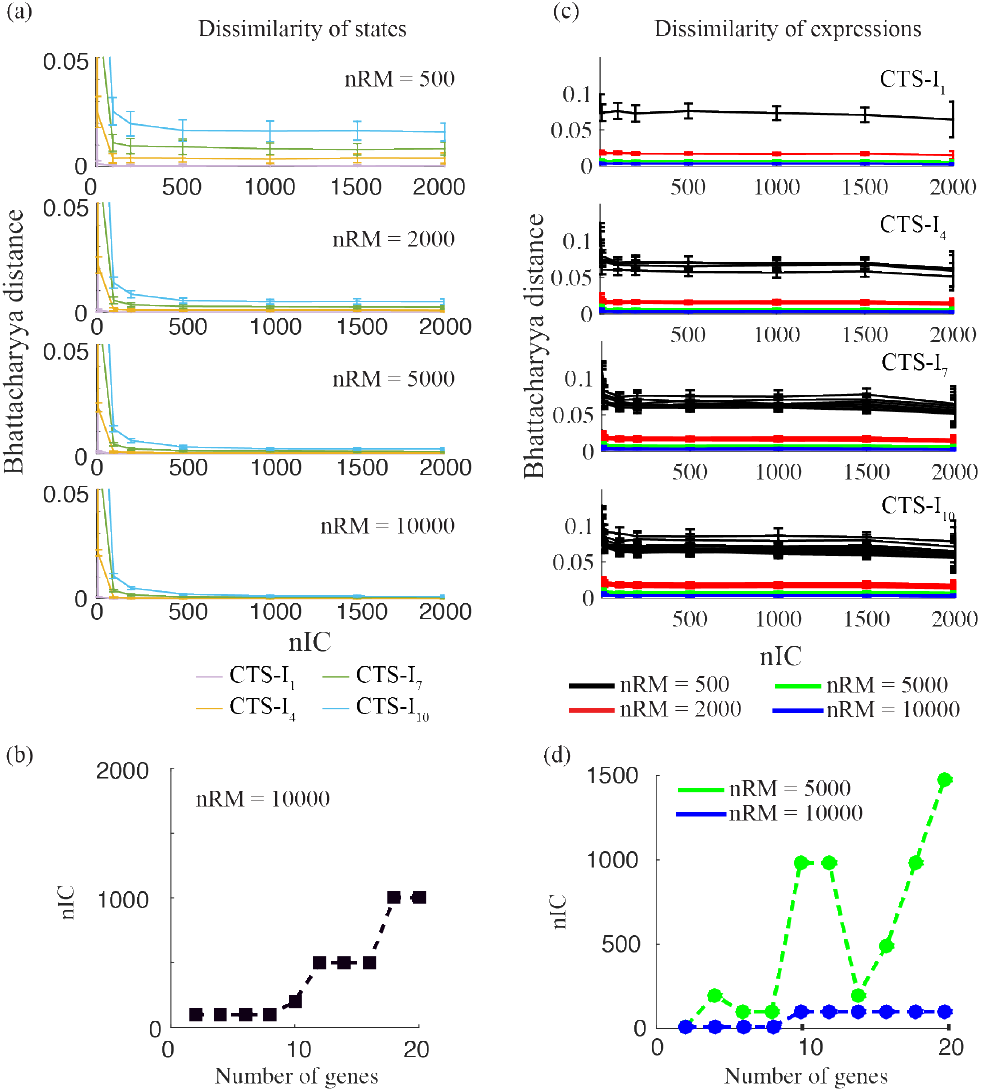
The effect of the number of initial conditions on the convergence of the RACIPE results. (a) For each coupled toggle-switch I (CTS-I) circuit (curves in different colors), the convergence is evaluated by the dissimilarity of states using different numbers of initial conditions (nIC in x-axis) and different numbers of RACIPE models (nRM in different panels). (b) The minimum nIC to get the converged distribution of the number of stables states when nRM equals to 10,000. Different points represent the CTS-I circuits of different sizes. The minimum nIC is selected if the decrease of the Bhattacharyya distance is smaller than the threshold (0.0005, see Fig. S2) when nIC increases. (c) For each CTS-I circuit, the convergence is alternatively evaluated by the dissimilarity of expressions of each gene. Only the Ai genes for each circuit are plotted (one line per gene) and colored differently for different nRMs. The dissimilarity is less sensitive to nIC, but is dramatically reduced with the increase of nRM. (d) The minimum nIC to get the converged distribution of expressions. The minimum nIC is selected if the decrease of the Bhattacharyya distance is smaller than the threshold (0.0005, see Fig. S5) when nIC increases. nRM needs to be larger than 5000 otherwise the distribution is not converged even with nIC is 2,000.

**Fig. 3.**
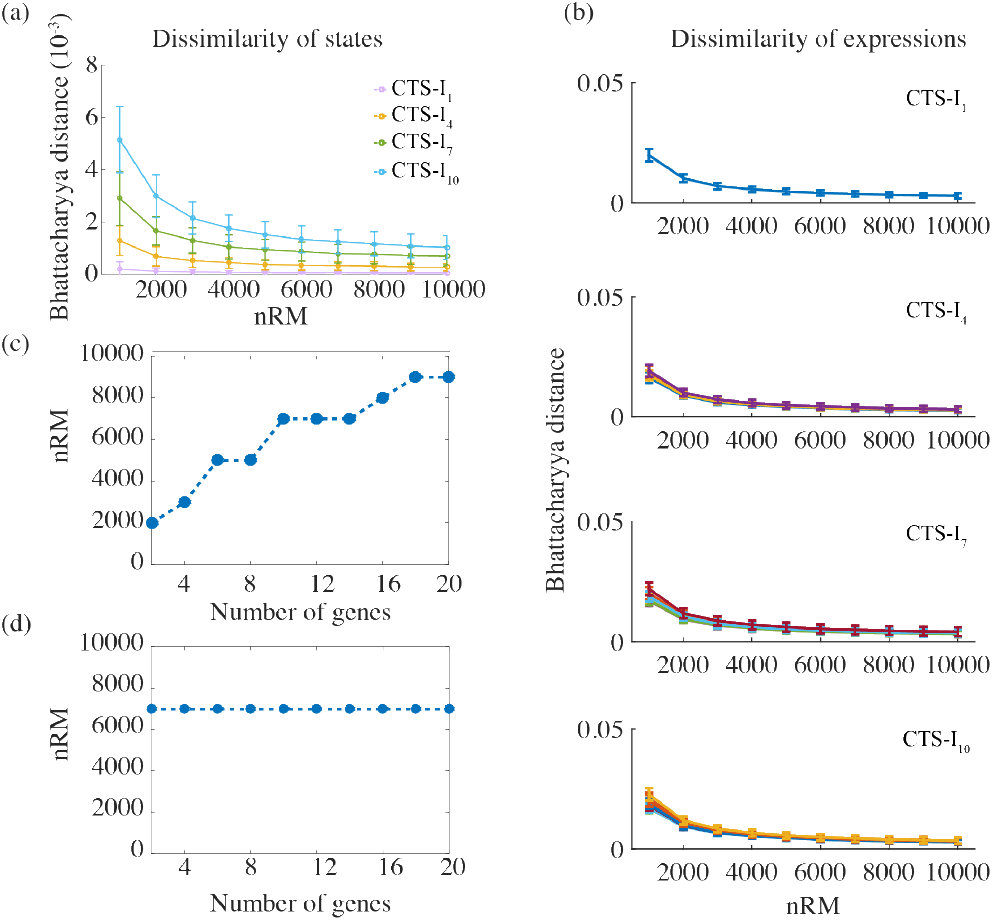
The effect of the number of RACIPE models on the convergence of the results. (a) The dissimilarity of states as a function of nRM when nIC is 1000. (b) The dissimilarity of expressions as a function of nRM when nIC is 1000. (c) The minimum nRM as the function of the number of genes in each circuit. (d) The minimum nRM to get the converged distribution of gene expressions.

For every choice of nIC and nRM, we repeat the RACIPE calculations for ten times for each circuit and measure the dissimilarity of the above-mentioned probability distributions by the Bhattacharyya distance 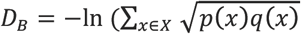, where *p* and *q* are two distributions. If the two distributions are exactly same, *D*_*B*_ equals to 0; if they are more different, *D*_*B*_ becomes larger.

To explore the effects of nRM on the distribution of the number of stable states, we repeat RACIPE on the circuit for ten times for a certain nRM, and calculate the distribution of the number of stable states for each replica. Then we compare the dissimilarity of the distributions (i.e. the dissimilarity of states) for different nRMs by calculating the average Bhattacharyya distances: 
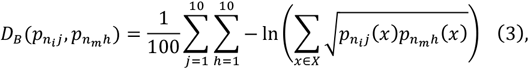
where *p*_*nij*_(*x*) stands for the probability of the circuit with *x* number of stable states for a random model for a replica *j* when nRM equals to *n_t_. n_m_* is the maximum nRM used in the test. Here we fix *n*_*m*_ to 10,000. Similarly, we can explore the effects of nRM on the distribution of gene expressions. Similar approach is used to measure of the effects of nIC.

As shown in Fig. 2a and Fig. S1-3, the dissimilarity of states decreases when more initial conditions are used. When nIC is larger than 500, RACIPE can effectively identify most stable steady states, except for some rare states (the probability to be observed is less than 1%). To get converged distribution of the number of stable states, the minimum required nIC increases with the size of the circuit (Fig. 2b and Fig. S3). Surprisingly, the convergence of the distribution of expressions seems to be less sensitive to nIC (Fig. 2c and Fig. S4-5), as similar results are obtained no matter how small or larger nICs are selected. As suggested from Fig. 2d, with more than 10,000 RACIPE models, 100 initial conditions are sufficient to get converged results.

However, nRM has a significant influence on the convergence of the simulation results. From Fig. 2a and Fig. S3, increasing nRM dramatically lowers the dissimilarity of states. Also, without enough RACIPE models, the distribution of expressions does not converge even when a large nIC is used (Fig. 2d). Furthermore, when nIC equals to 1000, both the dissimilarity of states and gene expressions decrease when nRM increases (Fig. 3a, 3b and Fig. S6-10). To get converged results for the distribution of states, the minimum required nRM again increases with the size of the circuit (Fig. 3c). However, the minimum required nRM to get the converged distribution of expressions is likely independent to the size of the circuit as long as it is more than 7,000 (Fig. 3d). Interestingly, when the dissimilarities of states for different circuits are scaled by the maximum number of stable states of the circuits, the curves of the dissimilarities for each circuit overlap with each other (Fig. S7b). The results suggest that the higher dissimilarity of a larger circuit is due to the higher complexity of the system.

### 3.3 Analysis of the RACIPE-generated Data

Once RACIPE generates, for each model, the kinetic parameters and the stable-state gene expressions, a variety of statistical methods can be applied to analyze the data from the ensemble of models. In the following, we will illustrate these analyses in the context of a coupled toggle-switch circuit (CTS-I_5_, with five toggle switches) (Fig. 4a). We generate 10,000 RACIPE models, each of which is simulated starting from 1,000 initial conditions. For each model, the maximum number of stable steady states is seven (Fig. S1); from 10000 RACIPE models, there are a total of 24,425 steady states. These states could be regarded as the gene expressions of cells in a system obeying these dynamics.

**Fig. 4.**
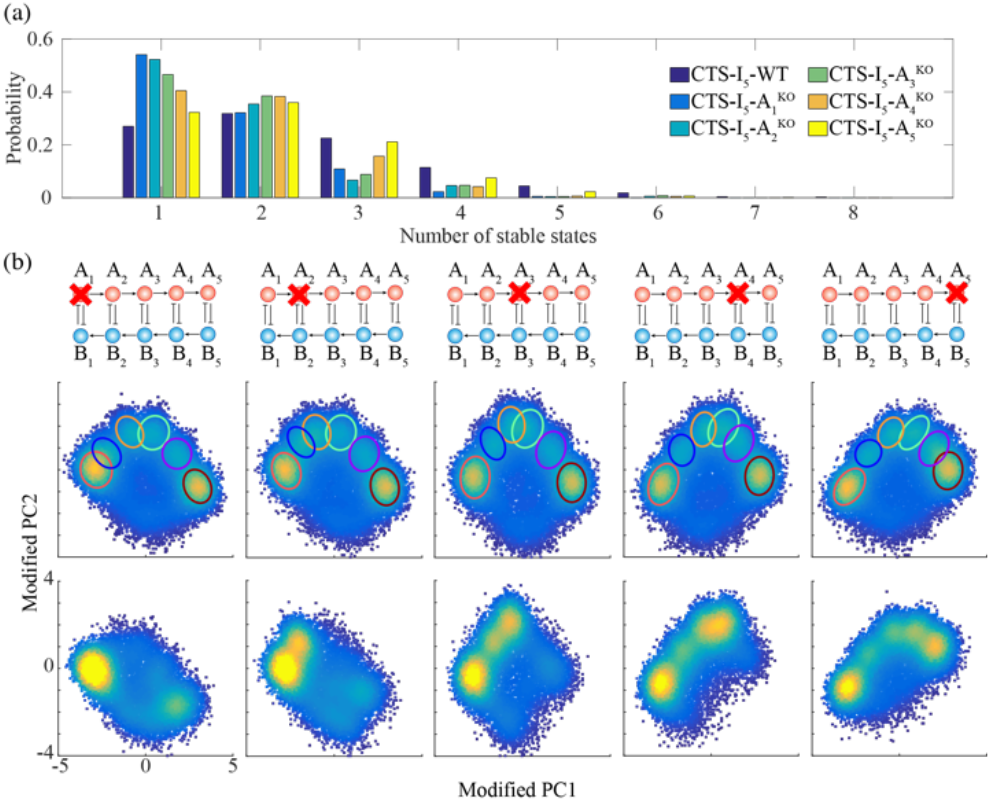
RACIPE identifies robust gene states of a coupled toggle-switch (CTS-I5) circuit. (a) Diagram of the CTS-I_5_ circuit. (b) Average linkage hierarchical clustering analysis of simulated gene expressions reveals six major clusters of distinct expression patterns. Each column corresponds to a gene, and each row corresponds to a stable steady state from a RACIPE model. (c). Histogram of the fraction of gene expressions in each cluster. The cutoff is selected at 5% (Red dash line). (d) 2D probability density map of the gene expressions projected on to the first two principal components. The six gene clusters are highlighted by the same colors as those in (b).

To analyze the simulated gene expression, RACIPE utilizes average linkage hierarchical clustering analysis (HCA) using Euclidean distance after normalization of the expressions (see SI 1.5-1.8 for details). From the heatmap (Fig. 4b), we observe six major clusters that have at least 5% fraction for each (Fig. 4c). The six major clusters, denoted by “gene state” below, are further confirmed by projecting all steady state solutions onto the first two principal components (PC1 and PC2) (Fig. 4d). From HCA, genes with similar functions are also grouped together. Strikingly, the gene expression patterns of the couple toggle-switch circuits, from the top to the bottom, correspond to a cascade of flips of the state of each toggle-switch motif (Fig. 4b). For instance, compared with gene state 2, gene state 5 has a flipped state in the fifth toggle-switch motif (A_5_ and B_5_).

Moreover, RACIPE can identify the roles of individual genes in the dynamic behaviors of the circuit by *in silico* gene knockouts, one gene at a time (Fig. 5 and Fig. S11). Knocking out gene A_1_ dramatically changes the probability distribution of the number of stable states and probability distribution of gene expressions, while knocking out gene A_5_ leads to a similar distribution of the number of stable states and only one gene state is missing. Therefore, we find that, for coupled toggle-switch circuits, the importance of A_i_ genes gradually decrease - A_1_ is the most critical one and A_5_ is the least important one. Similarity, the importance of B_i_ genes is in the reversed order. In addition, RACIPE can identify the significantly differentiated parameters between two states by the statistical analysis of model parameters (Fig. S12, see SI 1.9), which further helps to elucidate the functions of gene circuits.

**Fig. 5.**
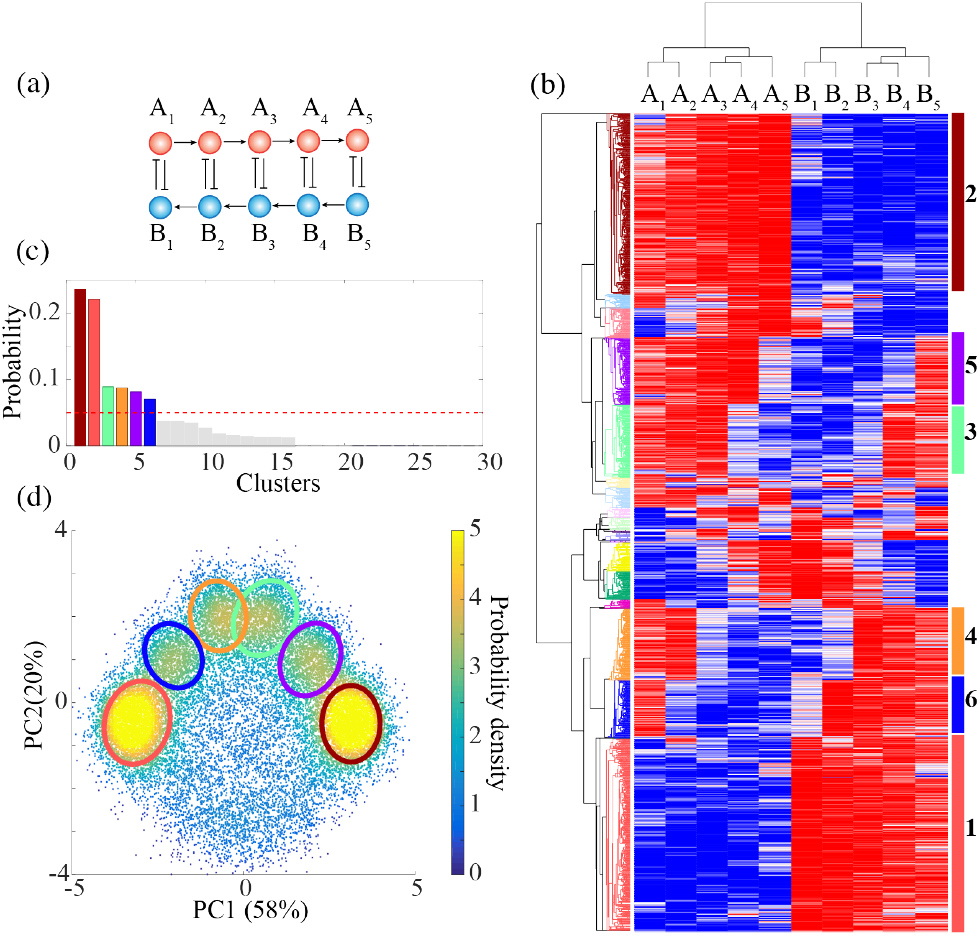
Perturbation analysis. (a) Probability distribution of the number of stable steady states of each model. Different colors represent the results of the standard RACIPE (CTS-I_5_-WT) and different knockout versions of RACIPE (CTS-I_5_-A_i_^KO^). (b) Probability density maps of the RACIPE gene expressions projected on to the first two principal components. Note, for the knockout cases, the principal components are modified to reflect the zero expressions for the corresponding genes (see SI for details).

### 3.4 Application to a B-lymphopoiesis Gene Circuit

The above example, while instructive, is only based on simple circuit motifs. To further evaluate the use of RACIPE, we analyze the properties of a gene regulatory circuit governing B-lymphopoiesis. This circuit was previously proposed by Salerno *et al.* (Salerno *et al.*, 2015) and analyzed mainly by traditional nonlinear dynamics methods, such as bifurcation analysis. Here we compare the RACIPE-generated gene expression data with microarray gene expression profiles for B cells from the previously published work by van Zelm *et al.* (van Zelm *et al.*, 2005).

B cells that develop in the bone marrow progress through the multipotent progenitor (characterized by CD34^+^/lin^−^), pro-B, pre-B-I and pre-B-II large, pre-B-II small and immature-B stages sequentially (van Zelm *et al.*, 2005). The regulatory circuitry for lineage specification of hematopoietic multipotent progenitors is still not well understood. To address this issue, Salerno *et al.* constructed a gene regulatory circuit (Fig. 6a) governing B-lymphopoiesis based on literature search and confirmed the important role of ZNF521 (zinc finger protein 521) and EBF1 (Early B-Cell Factor 1) during the specification of B cells from the multipotent progenitor stage (CD34^+^/lin^−^) to the pro-B stage (Salerno *et al.*, 2015). Here we apply RACIPE to the same gene network and study the predicted gene expression patterns and how they are associated with various stages during B cell development.

**Fig. 6.**
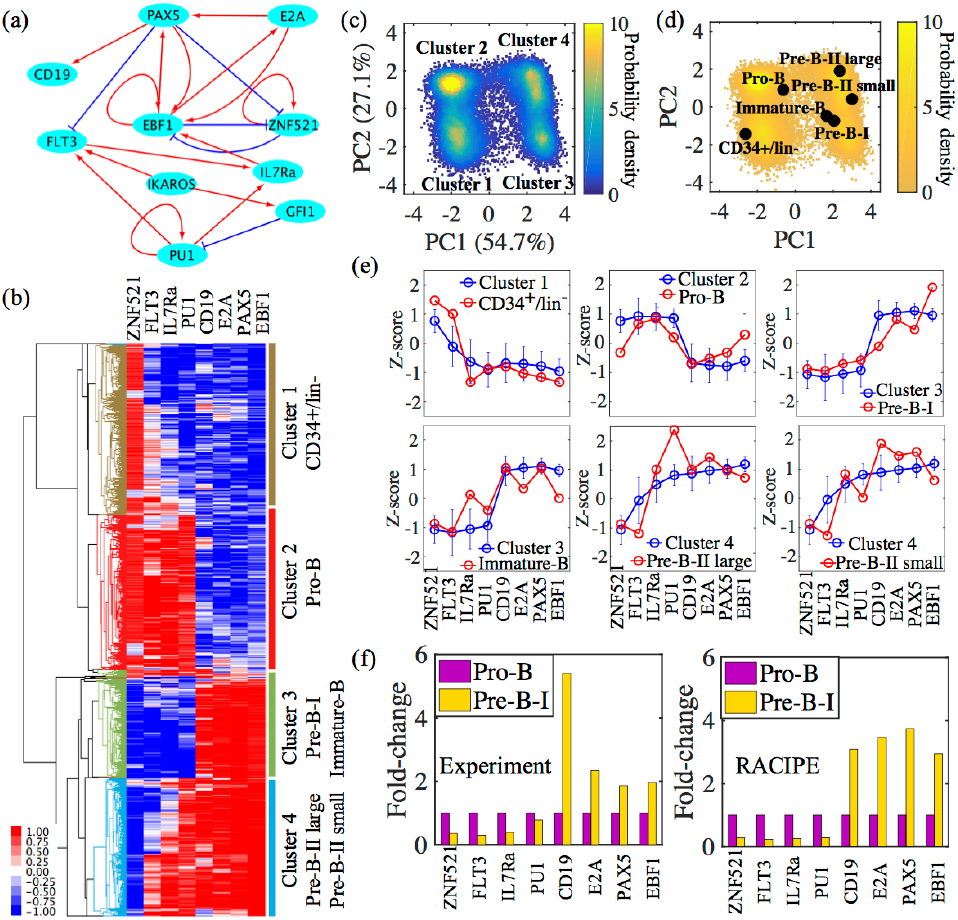
RAICPE identifies multiple gene expression states during B cell development. (a) A proposed gene regulatory circuit governing B-lymphopoiesis, adopted from (Salerno et al., 2015). The network consists of 10 transcription factors (TFs). Red arrows represent transcriptional activation and blue bar-headed arrows represent transcriptional inhibition. (b) Average linkage hierarchical clustering analysis of the gene expression data from all the RACIPE models using the Euclidean distance. Each column corresponds to a gene, and each row corresponds to a stable steady state. Four major gene states (clusters) are identified from the RACIPE analysis. (c) 2D probability density map of the RACIPE-predicted gene expression data projected on to the first two principal component axes. (d) The microarray expression profiling of different stages during B cell development (van Zelm et al., 2005) projected on to the same axes as shown in (c) (See SI 1.10). (e) Comparison between experimental gene expression of various stages with *in silico* clusters. Blue dots and red dots represent the Z-scores of genes from the RACIPE models and experiments, respectively. Error bar for each blue dot represents standard deviation of the RACIPE expression values. (f) Comparison between experimental gene expression fold-change from stage Pro-B to stage Pre-B-I with the computed fold-change by RACIPE.

Fig. S13 shows 10000 models are good enough to capture the robust behaviors of the gene network for B-lymphopoiesis. The stable steady states from all models form four major clusters, that correspond to the stages CD34^+^/lin^−^, pro-B, (pre-B-I, Immature-B) and (Pre-B-II large, small), respectively (Fig. 6b-d). We further compare the microarray gene expression profiles with data generated by RACIPE. Even through there is only one sample in each stage from (van Zelm *et al.*, 2005), the trend of the gene expression predicted by RACIPE agrees well with that from experiments, especially the comparison between cluster 1 and the CD34^+^/lin^−^ stage and that between cluster 3 and the Pre-B-I stage (Fig. 6e). From the hierarchical clustering analysis (Fig. 6b), we observe that there is a ‘switch-like’ change in the gene expression pattern from the stage pro-B to pre-B-I, as also shown in Fig. 6c. To test the prediction, we extract the microarray data of pro-B and pre-B-I and analyze the fold-change of the regulators in the circuit. Strikingly, the microarray data shows the down-regulation of TF ZNF521, FLT3, IL7Ra and PU.1 and up-regulation of CD19, E2A, PAX5 and EBF1, which validates the prediction from the RACIPE analysis (Fig. 6f). In summary, RACIPE is able to provide a rich source of information from the regulatory circuit of B-lymphopoiesis and potentially capture the gene expression features of various stages during B cell development.

Although we observe agreement between *in silico* clusters by RACIPE and microarray data of various stages in B cell development, we might not yet be able to generate all information regarding the paths of B cell development. The reasons are at least two-fold. First, the result by RACIPE is highly dependent on the topology of the gene circuit and there might be important genes/regulations missing in the current circuit due to insufficient knowledge from available data. Second, due to the very limited number of experimental samples, i.e., one in each stage, the comparison with clusters by RACIPE might be inaccurate. However, with even the limited information, RACIPE has been shown to capture the change of multiple master regulators across various stages during B cell development. Further study including construction of a more complete regulatory circuit for B cell development and measures of gene expression of more samples at various stages is needed to fully understand the state transitions of B cell progression.

## 4 Discussion

In this study, we introduced a new tool based on our recently developed computational algorithm, named *ra*ndom *ci*rcuit *pe*rturbation (RACIPE). Unlike traditional circuit modeling approaches that rely on a particular set of parameters, which might introduce incomplete or biased results, RACIPE generates an ensemble of models with random kinetic parameters, simulates the dynamics of these models by solving ODEs, and statistically analyzes the results. With this randomization approach, RACIPE can identify the most robust features of a gene circuit without the need to know detailed kinetic parameter values. In a sense, we convert a traditional non-linear dynamics problem into a statistical data analysis problem. The method has been implemented in C and will be freely available for academic use.

To better understand the performance of RACIPE, we particularly explored the effects of two important simulation parameters, the number of initial conditions (nIC) and the number of RACIPE models (nRM), on the convergence of the statistical analysis. Insufficient nIC and nRM may lead to inconsistent results in the repeats of the same simulation. Figs. 2 and 3 are good references for an initial guess of these parameters and users can always identify the optimal nIC and nRM with a similar analysis. From our tests, the time cost of RACIPE scales linearly with the total number of parameters used in the mathematical model, suggesting its potential use in analyzing large gene networks.

To illustrate the use of RACIPE, we applied it to a coupled toggle-switch (CTS-I_5_) circuit consisting of five toggle switches, a circuit that has an implication in coupled decision-making of multiple cell fates. From the RACIPE-generated expression data, we identified six major clusters by both HCA and PCA. In addition, we analyzed the role of each gene on circuit dynamics by *in silico* gene knockout (Fig. 5). To further show the predictive power of RACIPE, we applied the method on a published B-lymphopoiesis gene regulatory circuit. The gene expression patterns of various stages during B cell development can be efficiently captured by RACIPE. Notably, the fold-change of master regulators among the stage ‘Pro-B’ and ‘Pre-B-I’ predicted by RACIPE agrees well with that from the microarray data. These results show that RACIPE can not only reveal robust gene expression patterns, but also help uncover the design principle of the circuit.

The capability of RACPE in identifying circuit functions using a randomization approach reinforces the hypothesis that circuit dynamics are mainly determined by circuit topology (Klemm and Bornholdt, 2005) not by detailed kinetic parameters. Indeed, it is commonly believed that, through evolution, gene circuits of important pathways should be robustly designed to be functional (Li *et al.,* 2004) even in a dynamic and heterogeneous environment (Kaluza and Mikhailov, 2007). In RACIPE, we take the advantage of this feature to interrogate the robustness of a gene circuit by randomly perturbing all the kinetic parameters, from which we evaluate the most conserved properties.

Although we believe RACIPE have wide applications in systems biology, there are a few limitations of the current version. First, while all parameters are completely randomized to generate models, some of these models might not be realistic because some parameters are unlikely to be perturbed in cells, such as the number of binding sites. In these cases, incorporating relevant experimental evidences will improve the modeling. Second, RACIPE is unique in generating data of both gene expressions and model parameters. Although we have shown that the parameters in models from different gene state clusters are distinct (Fig. S11), further data analysis methods are needed to fully understand the roles of each parameter in circuit behavior. Third, current RACIPE only models regulatory circuits of transcription factors. However, the same approach can be extended to model biological pathways, which typically involves regulations of multiple types, such as protein-protein interactions and microRNA-mediated regulations. Fourth, we currently use deterministic ODE-based method to model the dynamics. Since gene expression noise has been shown to play crucial roles in circuit dynamics (Raser and O’Shea, 2005; Munsky *et al.,* 2012), it is important to extend RACIPE to stochastic analysis. Lastly, the quality of the circuit topology may dramatically impact the quality of RACIPE modeling. An accurate inference method for constructing gene circuits is especially important. Further improvements in these aspects will greatly improve the usability of this randomization-based approach and contribute to a better understanding of the operative mechanisms of gene regulatory circuits.

